# Microstructural pruning in human prefrontal cortex scaffolds its functional reorganization across development

**DOI:** 10.1101/2024.04.29.591716

**Authors:** Jewelia K. Yao, Zachariya Yazdani, Ruggaya Musa, Jesse Gomez

## Abstract

Neural representations in occipitotemporal cortex emerge during development in response to visual experience with ecological stimulus categories, such as faces or words. While similar category-selective representations have also been observed in the frontal lobe, how they emerge across development, whether current models of brain development extend to prefrontal cortex, and the extent to which such high-level representations are anatomically consistent across the lifespan is unknown. Through a combination of functional and quantitative MRI scans, we observe previously undescribed cortical folding patterns in human-specific inferior frontal cortex whose consistency reveals that childhood representations for visual object categories rearrange into stable adulthood patterns. This functional restructuring was distinct from occipitotemporal cortex where adult-like response patterns only scale in magnitude across development. The unique form of functional development in prefrontal cortex was accompanied by restructuring of cortical tissue properties: macromolecules are pruned across adolescence in prefrontal cortex while they proliferate in temporal cortex. These results suggest visual representations in distinct cortical lobes undergo distinct developmental trajectories, and that human-specific prefrontal cortex shows an especially protracted maturational process that necessitates late-stage tissue restructuring detectable in the living brain.

## Introduction

The visual ability to recognize ecological stimuli such as faces[1-3] and words[4,5] is a critical skill resulting from specialized neural representations in occipitotemporal visual cortex, which form in response to visual experience during childhood[2,3,6-8]. The consistency with which these regions emerge developmentally relative to cortical folding patterns[9-11] has made them useful for modeling[8,12] how the function and structure of human cortex matures to support behavioral learning[13,14]. While there has been much research charting how neural correlates underlying these visual abilities develop in sensory cortex[8,12,13,15], how higher-order visual representations in prefrontal cortex involved in working memory and attention[16] emerge during childhood remain relatively opaque.

Such representations lie largely within ventrolateral prefrontal cortex (VLPFC)[16,17], whose visual responses have likely remained uncharted in children for two reasons. Firstly, multimodal pediatric neuroimaging is difficult, especially when attempting to measure such high-level representations. Secondly, a strong neuroanatomical model linking cortical folds with functional representations does not exist for prefrontal representations as it does for occipitotemporal visual cortex[10,18-22], leaving uncertainty in whether observed responses are anatomically similar or distinct over development. The folding of the cortical sheet is a hallmark feature of the human brain[23,24], with growing evidence suggesting that cortical folds are predictive landmarks of functional representations within cortex well beyond primary sensory representations[10,20,25]. Many smaller folds, known as tertiary sulci, can only be observed in humans and have been increasingly shown to underlie human-specific cognitive abilities. Importantly, this has been demonstrated in human prefrontal cortex, with variability in sulcal patterns of the superior and middle frontal sulci predicting differences in complex reasoning abilities[26-29]. However, visually-responsive representations for faces[17,30,31] and words[32-34] are located more posterior and ventrally within the precentral (PCS) and inferior frontal sulci (IFS), where such a model has not yet been described. Demonstrating anatomically-predictable visual representations in prefrontal cortex would extend our understanding of structure-function coupling beyond classic sensory regions into the apex of association cortex.

The cortex surrounding the IFS is evolutionarily recent, specific to humans and great apes [35], thereby representing an opportunity to ask how cortex that is novel on an evolutionary scale may develop in a functionally and structurally unique manner. Current models of visual brain development linking experience to functional and structural maturation have been based on ventral temporal cortex (VTC), where regions involved in face and word processing show relatively linear growth across development[2,5,14], coupled with growth in underlying cortical tissue structures[36]. However, synaptic work primarily from non-human primates has shown that higher-level prefrontal cortex undergoes more protracted[37,38] and sometimes qualitatively unique[39] development. Thus, whether or not developmental models of functional and structural brain development established in posterior visual cortex are universally true of all cortex in the human brain remains unknown.

Here, we combine multimodal neuroimaging of the human brain, measuring visual responses to ecological visual categories with functional MRI and fine-scale tissue properties through recent advances in quantitative MRI sensitive to the precise quantity and composition of neural tissue in the living brain. We scan both children and adults to investigate (i) to what extent ventrolateral prefrontal cortex demonstrates category-selective response to ecological visual stimuli, (ii) how these representations emerge over development, (iii) identify whether visual responses are consistent relative to cortical folding patterns, and (iv) establish whether functional development in prefrontal cortex necessitates changes in fine-scale tissue properties in the underlying cortical sheet. We include measurements from ventral temporal cortex to further determine whether two visually-responsive brain regions located in separate lobes show similar or distinct developmental trajectories. We find that despite clear and adult-like visual patterns in temporal cortex, responses in prefrontal cortex show especially protracted development, with childhood visual responses reorganizing into stable adulthood patterns. Prefrontal visual responses in adulthood seem to follow sulcal pleating patterns which have yet to be described in human neuroanatomy. These distinct functional maturation patterns are accompanied by equally distinct changes in fine-scale tissue properties, with prefrontal cortex demonstrating late adolescent pruning of both myelinated and non-myelinated structures, which we validate with developmental gene expression analyses. Our results suggest visual representations in different cortical lobes undergo distinct trajectories of functional and structural development, with prefrontal cortex showing unique, late-stage pruning of cortical tissue detectable in the living brain.

## Results

Although visual responses in VLPFC have only been reported in adults, we hypothesize PFC will show visual responses in children, albeit to a lesser-developed extent. To quantify this, 25 children ages 5 to 11 years old (mean = 8.92 ± 2.18, 14 females) and 20 adults ages 22 to 26 years old (mean = 23.55 ± 1.23, 15 females) completed structural MRI to reconstruct the cortical surface, and functional MRI while performing a visual category localizer containing visual stimuli of faces, words, bodies, objects, and places. To assess if prefrontal cortex is generally responsive to visual stimuli, we define a broad region of interest (ROI) encompassing the PCS and IFS where category-selective regions have been previously reported in adults[16]. From this swath of cortex, we can quantify the surface area selectively responsive to different visual stimuli, and further quantify if PFC, like occipitotemporal cortex, demonstrates previously observed asymmetries in the representation of certain stimuli. While hemispheric lateralization has been reported for faces and words in ventral temporal cortex[40-44], the extent these hemispheric asymmetries exists in frontal cortex beyond the well-known left-lateralization of language[45] and how they further emerge across development is not clear. In PFC, we find that PFC activity is indeed driven by visual stimuli, with children and adults both showing selective responses to all stimulus categories (**Fig 1A**). The left hemisphere responses are dominated by word representations, which increase by 38% from childhood to adulthood. In the right hemisphere, word responses appear to decrease by 16.5% while responses to faces increase by 89%. In both hemispheres, the representation of objects seems to be pruned back with development, decreasing by 45% in the left and 47% in the right hemisphere. An ANOVA in left VLPFC with factors of age-group and category reveals no significant interaction of age and category (F(1,4)=0.96, p=0.43), while an ANOVA in right VLPFC finds a significant age-category interaction (F(1,4)=2.7, p<0.03). This differential development leads to an uneven distribution, or lateralization, of category representations across hemispheres (**Fig 1B**).

**Figure 1.**
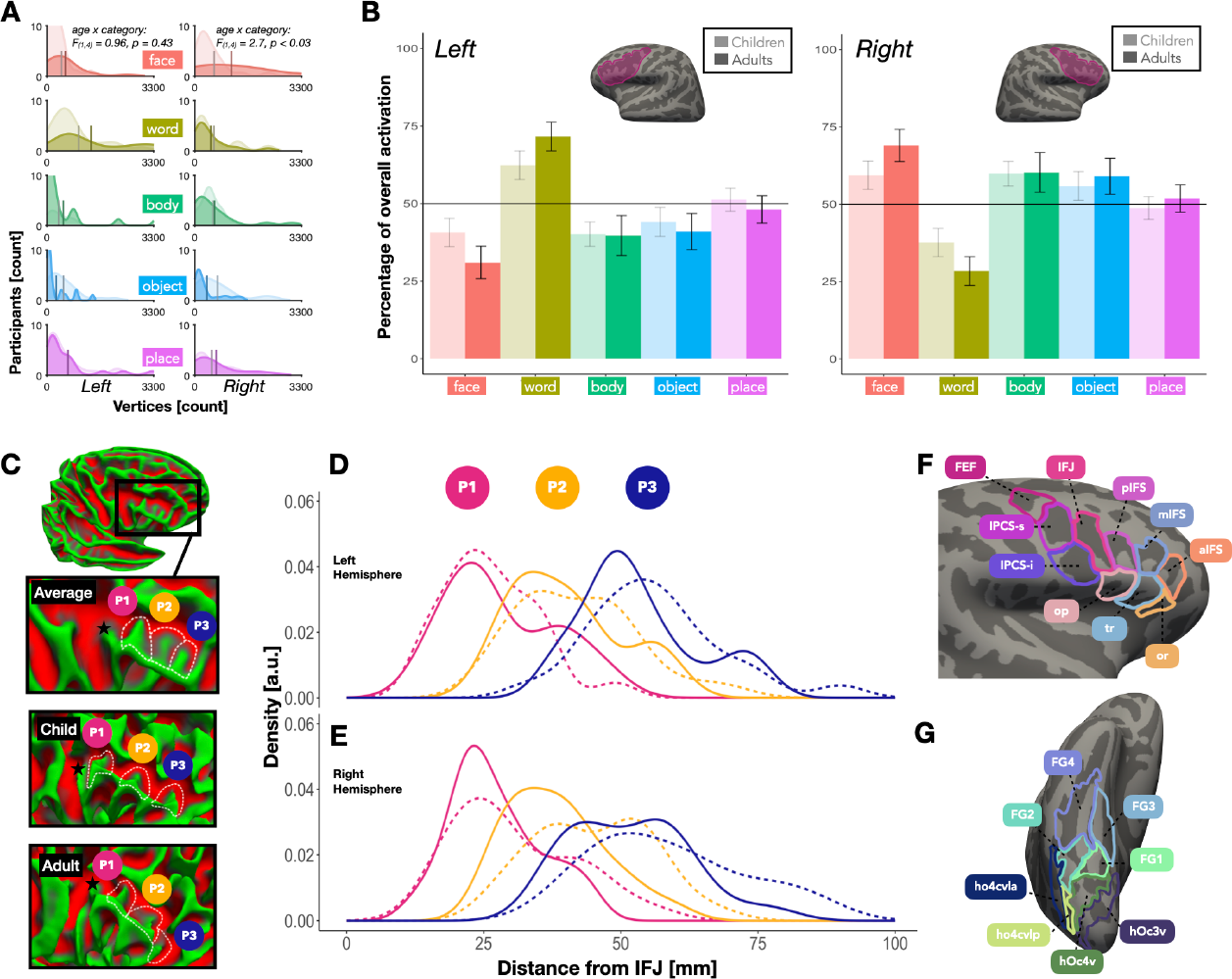
Ventrolateral prefrontal cortex shows both visual responses and novel folding patterns across development. **(A)** Histograms of surface area in VLPFC showing selective responses to each category (contrast threshold of t-values > 2.5) in the left and right hemispheres. Histogram height corresponds to the number of participants. Children are light colors, adults dark colors. **(B)** Lateralization of each category in children and adults. Inset: Outline of regions in VLPFC (pink) based on boundaries defined in Levy and Wagner (2011) and Bugatus et al. (2017). Left: Comparison of activation in the left hemisphere for faces (red), words (yellow), bodies (green), objects (blue), and places (purple) in children and adults. Bars above the 50% line indicate lateralization to the left hemisphere. Error bars denote standard error of the mean. Right: Same as left panel, but for the right hemisphere. **(C)** Examples of IFS pleats on surfaces of the fsaverage, an example child, and an example adult. IFS-P1 (pink), IFS-P2 (yellow), IFS-P3 (blue) are demarcated by white dotted lines. Black stars represent the location of the inferior frontal junction (IFJ). **(D)** Histogram of distances of the central vertex of IFS-P1, IFS-P2, and IFS-P3 from the IFJ (black star) in children (dotted lines), and adults (solid lines) in the left hemisphere. **(E)** Same as B but for the right hemisphere. **(F)** Parcellation of VLPFC into 9 ROIs derived from the precentral sulcus and IFS pleats, and areas of the inferior frontal gyrus. IFJ: inferior frontal junction, pIFS: posterior inferior frontal sulcus, mIFS: middle inferior frontal sulcus, aIFS: anterior inferior frontal sulcus, FEF: frontal eye fields, IPCS-s: superior component of inferior precentral sulcus, IPCS-i: inferior component of inferior precentral sulcus, op: pars opercularis, tr: pars triangular, or: pars orbitalis. **(G)** Parcellation of VTC into 8 ROIs from Grill-Spector and Weiner (2014).

To quantify changes in hemispheric lateralization, a multiple linear regression revealed a hemisphere by category effect whereby individual categories, specifically words and faces, are lateralized to either the left or right hemisphere (F(23, 516) = 5.83, p < 0.0001). To further test for lateralization for faces and words, we calculated the laterality index (LI) – the percentage of voxels with t-values greater than 2.5 that are lateralized to one hemisphere rather than the other – in VLPFC for each participant and then performed a one-sample t-test to determine if the group mean LI is significantly different from zero. The results of the paired-sample t-tests indicate that both the LI for faces (t (1,44) = 3.94, p = 0.0003) and the LI for words (t (1,44) = 4.97, p = 1.05e-05) were significantly different from zero, indicating prefrontal lateralization for the processing of faces and words. In particular, faces are lateralized to the right hemisphere and words are lateralized to the left hemisphere in both childhood and adulthood. Bodies and objects are also more right lateralized (bodies: t (1,44) = 2.83, p = 0.007, objects: t(1,44) = 2.01, p = 0.05) but place activations are found more evenly across both hemispheres (t(1,44) = 0.05, p = 0.96). For faces and words, development entails trending increases in lateralization, as word responses are pruned in the right and increase in the left (RH: Children= 37.62, Adults= 28.38; LH: Children= 62.38, Adults= 71.62; t(1, 42.25) = 1.42, p = 0.16) with face responses showing the opposite pattern (RH: Children= 59.36, Adults= 68.99; LH: Children= 40.64, Adults= 31.01; t(1, 40.32) = 1.39, p = 0.17). For bodies, objects, and places, there are no significant changes in lateralization from childhood into adulthood (bodies: t (1,32.32) = 0.06, p = 0.96, places: t (1, 39.58) = 0.55, p = 0.59, objects: t (1, 38.47) = 0.42, p = 0.68).

While prefrontal cortex demonstrates visual responses to categories that vary across hemisphere, how might the functional topography within VLPFC change across childhood? Do topological changes in prefrontal responses simply mirror those of ventral temporal development, or do different lobes show different patterns of maturation? To answer these questions, and explore whether functional representations in prefrontal cortex are associated with cortical folding, we can take advantage of previous research in occipitotemporal cortex showing that borders between functionally-distinct regions follow cortical folds, including primary, secondary, and tertiary sulci[10]. Defining an anatomically-based parcellation will allow us to quantify the location and spatial distribution of visual responses consistently across participants. Such a map of functionally-relevant cortical folds has yet to be produced in inferior prefrontal cortex. One unique aspect of our region of interest is that all visual responses are located within major sulci (PCS or IFS), and therefore any neuroanatomical model would have to describe features within a sulcus. Such a model was first proposed in 1885[46], where Louis Pierre Gratiolet described that some larger sulci in the primate brain showed consistent folding patterns along their sulcal walls. These intra-sulcal folds were deemed *plis de passage* or “passage pleats”, later translated as “annectant gyri”[24,47,48]. We find that both the PCS and IFS not only demonstrate this pleating phenomenon, but do so consistently across participants (**Fig 1C**). The PCS is historically divided into superior and inferior components[49], separated by what we identify here as annectant gyri, one located just posterior to middle frontal gyrus, and a second annectant gyrus which further divides the IPCS into inferior and superior halves (**Fig 1F**).

Within the IFS, we find three novel pleats formed by three annectant gyri separated by smaller sulci within the IFS. Calculating the geodesic distance between the central vertex of each pleat’s sulcal bed and a vertex located at the most posterior point of the IFS in each individual (e.g., the junction with the precentral sulcus), we find that the first pleat (IFS-P1) is consistently found closest to the beginning of the IFS; Child means: RH = 30.52 ± 11 mm, LH = 26.29 ± 8.47 mm, Adult means: RH = 27.20 ± 8.27 mm, LH = 28.35 ± 9.94 mm. The second pleat IFS-P2 sits above the anterior half of the *pars opercularis* (*op*); Child means: RH = 45.04 ± 10.51 mm, LH = 41.96 ± 10.78 mm, Adult means: RH = 28.25 ± 8.90 mm, LH = 40.10 ± 9.82 mm). IFS-P3 borders the posterior half of the *pars triangularis* (*tr*); Child means: RH = 58.84 ± 13.89 mm, LH = 56.21 ± 12.36 mm, Adult means: RH = 51.65 ± 10.72 mm, LH = 53.25 ±10.72 mm, and IFS-P3 superiorly spans the anterior half of the *tr*. Across individuals, histograms of distances of these gyri from the most posterior point of the IFS follow roughly normal distributions, with P3 showing a slight rightward skew (**Fig 2D-E**). For all three pleats in the left hemisphere, two-sample t-tests comparing children and adults reveal no significant differences in pleat distance from the IFJ (P1: p = 0.47, P2: p = 0.55, P3: p = 0.40). However, this was not the case in the right hemisphere. While there is no difference in distance for P1 (t (1,42.87) = 1.16, p = 0.25), children and adults show significant differences in the location of P2 (t (1,42.84) = 2.34, p = 0.02) and trending differences in the location of P3 (t (1,42.96) = 2.34, p = 0.06) in the right hemisphere.

**Figure 2.**
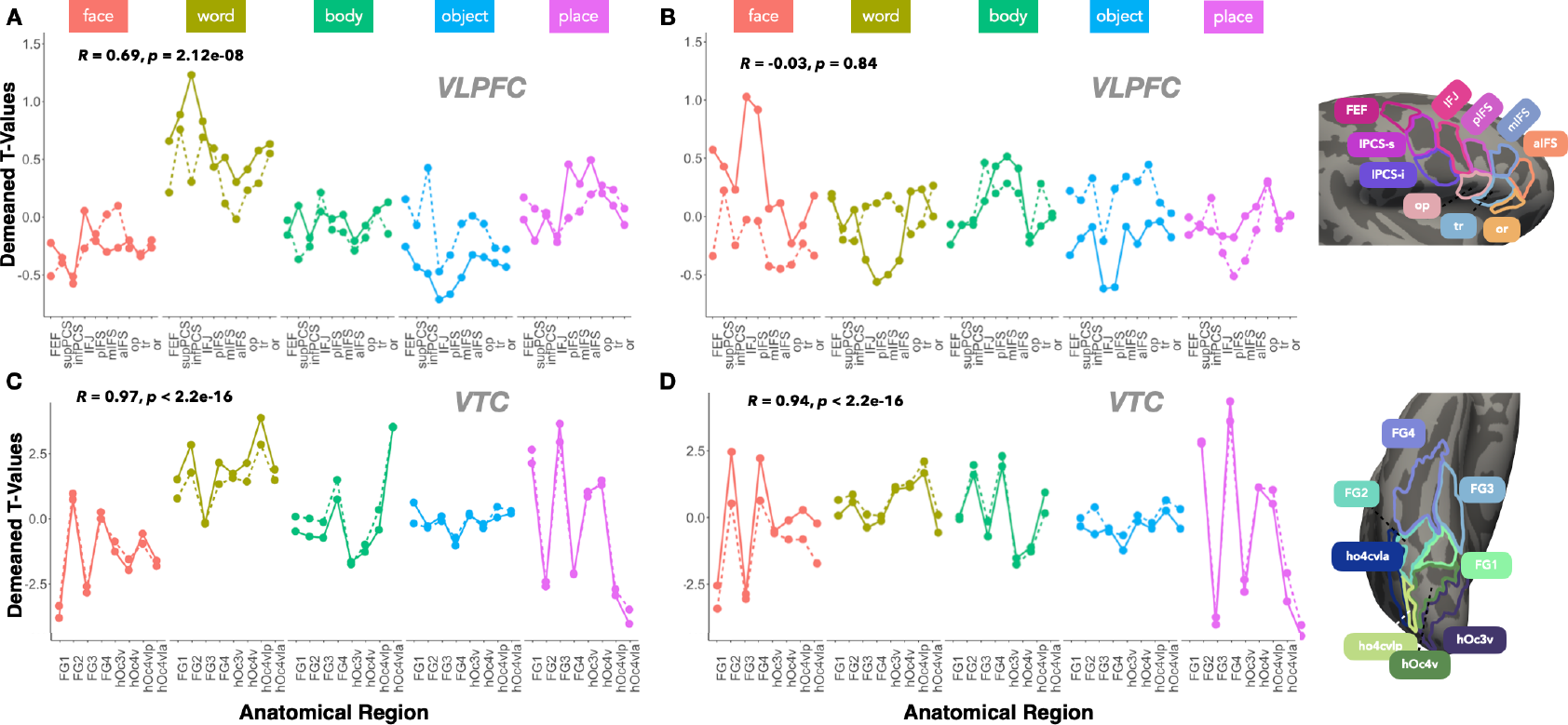
Broad category response patterns develop more in VLPFC than VTC, and more so in the right hemisphere. **(A)** Lineplots of average t-values, demeaned by the average response across all categories and ROIs for children and adults separately, to faces (red), words (yellow), bodies (green), objects (blue), and places (purple) in each of the 9 left hemisphere VLPFC ROIs. Dotted lines represent child data, and solid lines represent adult data. Pearson correlations between average adult and child responses across all categories and ROIs shown in the top left corner of each plot. **(B)** Same as panel A but for the right hemisphere. **(C)** Same as panel A but for the 8 left hemisphere VTC ROIs. **(D)** Same as panel C but for the right hemisphere.

Given their qualitative presence in all participants, the annectant folds of the IFS, together with annectant gyri of the PCS (**Fig 1C**) and the secondary sulci inferior to the IFS which form part of the pari-sylvan network[50-52] (*op, tr*, and the pars orbitalis *or*) represent stable neuroanatomical landmarks through which we create a parcellation of visually-responsive prefrontal cortex (**Fig 1F**). To compare prefrontal visual responses to those in classic visual cortex, we used 8 previously defined cytoarchitectonic regions located within occipitotemporal cortex whose boundaries similarly follow cortical folds[53,54]. We referenced FG1, FG2, FG3, FG4, hOc3v, hOc4v, hO3cvla, and hO4cvlp as VTC ROIs, which were projected from the fsaverage to every individual (**Fig 1G**). These occipitoemporal regions encompass the relevant cortex in which category-selective responses to our stimuli of interest are located. Does the topography of prefrontal responses remain stable and become scaled into adulthood, or does functional representation change its spatial pattern across development? Finer-grained analysis reveals that face, word, body, object, and place responses vary within and across ROIs. As expected, responses within VTC to each category are not uniform across the 8 ROIs (**Fig 2**). For example, in adults, responses to faces in VTC are highest in FG2 (lateral Fusiform) and lowest in hOc4vla (**Fig 2C-D**). Conversely, within an ROI, responses to different categories are not uniform; in FG1, there are high responses to places and low responses to faces. We mapped responses to individual categories within each ROI separately for children and adults. In VTC, we find that from childhood to adulthood, responses to categories show modest scaling but maintain their activity pattern across ROIs. For example, face responses in right VTC and word responses in left VTC scale positively with development. Overall, the patterns of responses between children and adults look very similar, so similar that the category-response pattern across ROIs is nearly perfectly correlated between children and adults in both hemispheres (RH: Pearson’s r = 0.94, p < 2.2e-16; LH: r = 0.97, p < 2.2e-16; **Fig 3A-B**).

**Figure 3.**
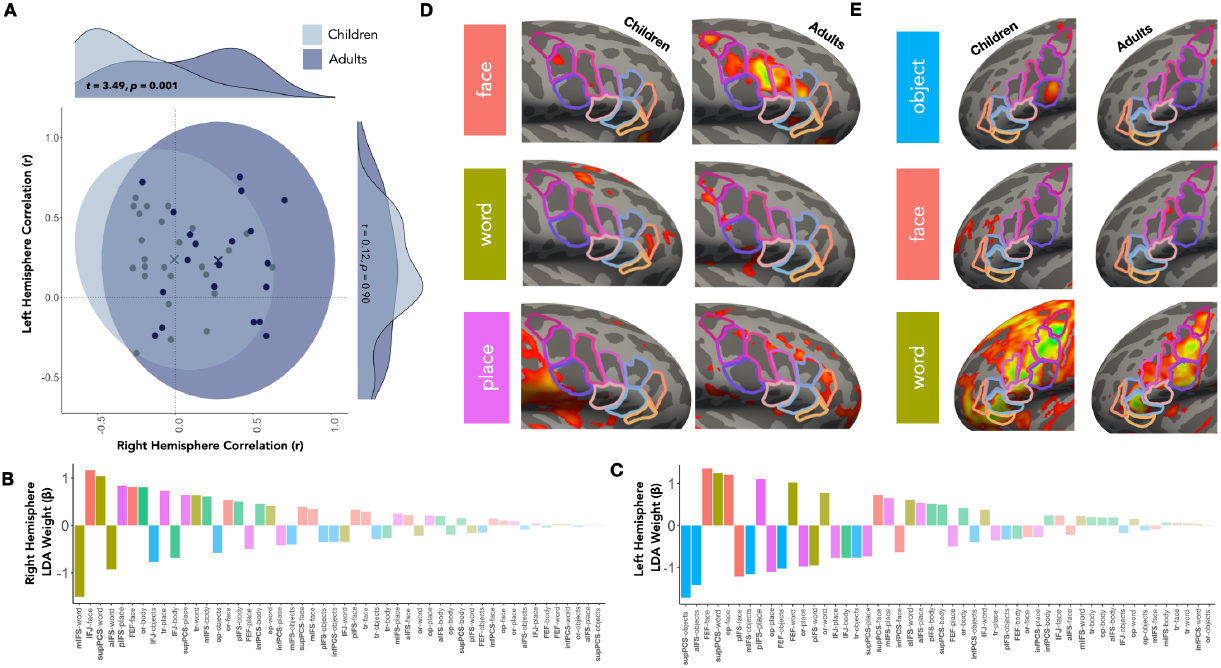
Differential development of category responses based on hemisphere, category, and region of interest. **(A)** Correlations of each child (light blue dots) and each adult (dark blue dots) to the adult average in the right hemisphere (x-axis) and left hemisphere (y-axis). Dark blue circle encompasses adults, with the dark blue x marking the adult centroid. Light blue circle encompasses child data, light blue x marks the child centroid. Distributions of correlation values and t-statistics comparing groups are shown on each axis. **(B)** Beta weights for each of the ROI-category coefficients in the linear discriminant analysis (LDA) separating children and adults in the right hemisphere. Strength of beta weight represents magnitude of change, and sign of beta weight represents scalar of change for faces (red), word (yellow), bodies (green), objects (blue), places (purple) in each ROI from childhood into adulthood. **(C)** Same as panel B for the left hemisphere. **(D-E)** Average cortical surface maps of face, word, place and object responses. Selectivity maps (contrast of preferred > non-preferred responses, thresholded at t-values > 2.5) were averaged across participants on the fsaverage brain.

As in VTC, we find that there are differential responses to categories across ROIs in VLPFC. However, the topographic changes in functional responses during development appear distinct from those of occipitotemporal cortex. In the right hemisphere VLPFC, we find that patterns seem to be resculpted across development whereby responses to categories might increase or decrease depending on the ROI and category (**Fig 2**). Interestingly, response patterns between children and adults are so distinct in the right hemisphere that they are not correlated across ROIs (R = -0.03, p = 0.84; **Fig 2B**). In left hemisphere VLPFC, the response pattern between children and adults are more similar, showing significant correlation (R= 0.69, p = 2.23-08; **Fig 2A**) suggesting that response patterns between age-groups are somewhat shared but there remains evidence of resculpting. For example, the ROI with the highest response to faces or objects in childhood is not the same ROI as in adulthood. Thus while both hemispheres show evidence of functional resculpting, the right hemisphere shows this maturation pattern more distinctly than the left. To further investigate the unique development observed in the right hemisphere, we conducted a correlation analysis between each individual’s response to each category within an ROI and the average response of adults for each category (**Fig 3A**). For adults, a leave-one-out approach was taken to avoid using an individual’s data in both the test and training data. In the right hemisphere, a Welch’s two-sample t-test comparing Fisher’s z-transformed correlations of children and adults to the adult average revealed a significant difference between group means (t (1, 36.49) = 3.49, p = 0.0001; **Fig 3A**). Specifically, adults exhibited higher correlation (M = 0.31) to the adult average than the children (M = - 0. 002). The same correlation analysis run in the left hemisphere did not reveal statistically significant differences between correlations of children and adult responses with the standard response (Madults= 0.27, Mkids= 0.26, t(1, 34.71) = 0.12, p = 0.90; **Fig 3A**). Together, this highlights unique developmental differences in responses for right hemisphere VLPFC.

To get a better sense of the representational changes driving prefrontal functional development, we ran a linear discriminant analysis (LDA) for each hemisphere to determine the categories and regions within VLPFC that best predict group differences. That is, the LDA identifies a linear combination of predictor variables that separate children and adults based on their response patterns. The 45 input variables corresponded to a particular category in a particular ROI (i.e. mIFS-word or IFJ-face), the strength of beta weights for the predictors indicates the magnitude. Examining the top five predictors in the right hemisphere LDA reveals that the main drivers of change were words (*β*rh.mIFS-word = -1.51, *β*rh.supPCS-word = 1.04, *β*rh.aIFS-word = -0.92), faces (*β*rh.IFJ-face = 1.17), and places (*β*rh.pIFS-place= 0.84; **Fig 3B**). Development in the left hemisphere was primarily driven by changes in responses to objects (*β*lh.supPCS-objects = 1.71, *β*rh.aIFS-objects = -1.41) as well as faces (*β*rh.FEF-face = 1.35, *β*rh.op-face = 1.21) and words (*β*lh.supPCS-words = 1.24; **Fig 3C**). Average contrast maps for the top three categories showing high LDA weights in each hemisphere are shown in **Figure 3D-E**. Notably, these changes in category responses are specific to anatomically-defined ROIs. While the beta-weights were comparable in magnitude across hemispheres, the accuracy of the left hemisphere LDA model in characterizing each individual as either a child or adult (44%) was lower than the accuracy of the right hemisphere model (78%). This mirrors the correlation analysis where right hemisphere response patterns are more discriminable between children and adults.

In previous work examining ventral temporal cortex[36], developmental increases in visual responses were hallmarked by the proliferation of fine-scale tissue compartments such as myelin and neuropil. If prefrontal responses show both increases and decreases across development, will the development of the underlying fine-scale tissue structures also show non-linear development? To quantify these micro-scale changes in cortical tissue, and complement the mesoscale anatomical measures of cortical folding presented earlier, we employed a quantitative MRI dataset with N = 82 individuals across the lifespan from 5 to 54 years of age[55]. The qMRI data provides brain-wide maps sensitive to tissue properties likely to develop[56]. Here, we examine changes in R1 (1/T1 where T1 is the proton relaxation time) which is a measure reflective of tissue composition and is especially sensitive to myelin. We also examine maps of macromolecular tissue volume (MTV), quantifying the volume (milliliters) of tissue within a voxel. We hypothesize that the differential functional development in VTC (scaling) and VLPFC (restructuring) necessitates distinct structural development within the underlying cortical tissue. To test this, we average tissue properties across cortical voxels within the larger VTC and VLPFC regions for each individual, and in each hemisphere.

In VTC, we find results consistent with previous work[36]. Gradual increases in R1 (**Fig 4A**) and MTV (**Fig 4B**) from childhood to older adulthood suggest water in the cortex is being continuously displaced by macromolecules, including myelin. This relatively linear growth in tissue across the lifespan does not appear to be shared with VLPFC, particularly within the right hemisphere. In right prefrontal cortex, while MTV and R1 values increase from childhood to teenage years, there appears to be a dip in trajectory in late adolescence, with young adults showing less cortical tissue microstructure compared to teens (**Fig 4A-B**). This dip is not visible in left VLPFC. To quantify this adolescent reduction in the right hemisphere, we ran an ANOVA comparing teens and young adults across factors of cortical lobe (temporal and frontal), which revealed a significant interaction (F(1,29) = 6.18, p < 0.001), supporting the notion that temporal and prefrontal cortex show distinct trajectories of tissue growth. An additional ANOVA showed an interaction of age-group by hemisphere (F(1,29)= 5.64, p = 0.02) when comparing left and right VLPFC, demonstrating the two prefrontal hemispheres undergo distinct tissue maturation trajectories as well. Thus, young adults appear to demonstrate some pruning or tissue loss in the right hemisphere, yielding prefrontal cortex tissue values similar to those in childhood. To replicate this observation, the same children and young adults who completed the aforementioned fMRI experiments were also invited to complete quantitative imaging. Because these age groups overlap with the children and young adult age groups from the larger qMRI dataset, we expect *a priori* to find no difference in tissue values in prefrontal cortex. In the right hemisphere, we verify this prediction, whereby children (N = 25) and adults (N = 20) show statistically similar R1 values (t(1,40.23) = -0.66, p = 0.51) and MTV values (t(1, 38.68) = -1.54, p = 0.13). In ventral temporal cortex, the same participants demonstrated significant tissue proliferation (increasing R1 values) from childhood to adulthood[36], as also observed in the present data (**Fig 4A-B**). Thus, two separate datasets confirm that right prefrontal tissue shows a unique developmental trajectory, one that involves fleeting loss of tissue microstructures in late adolescence to young adulthood.

**Figure 4.**
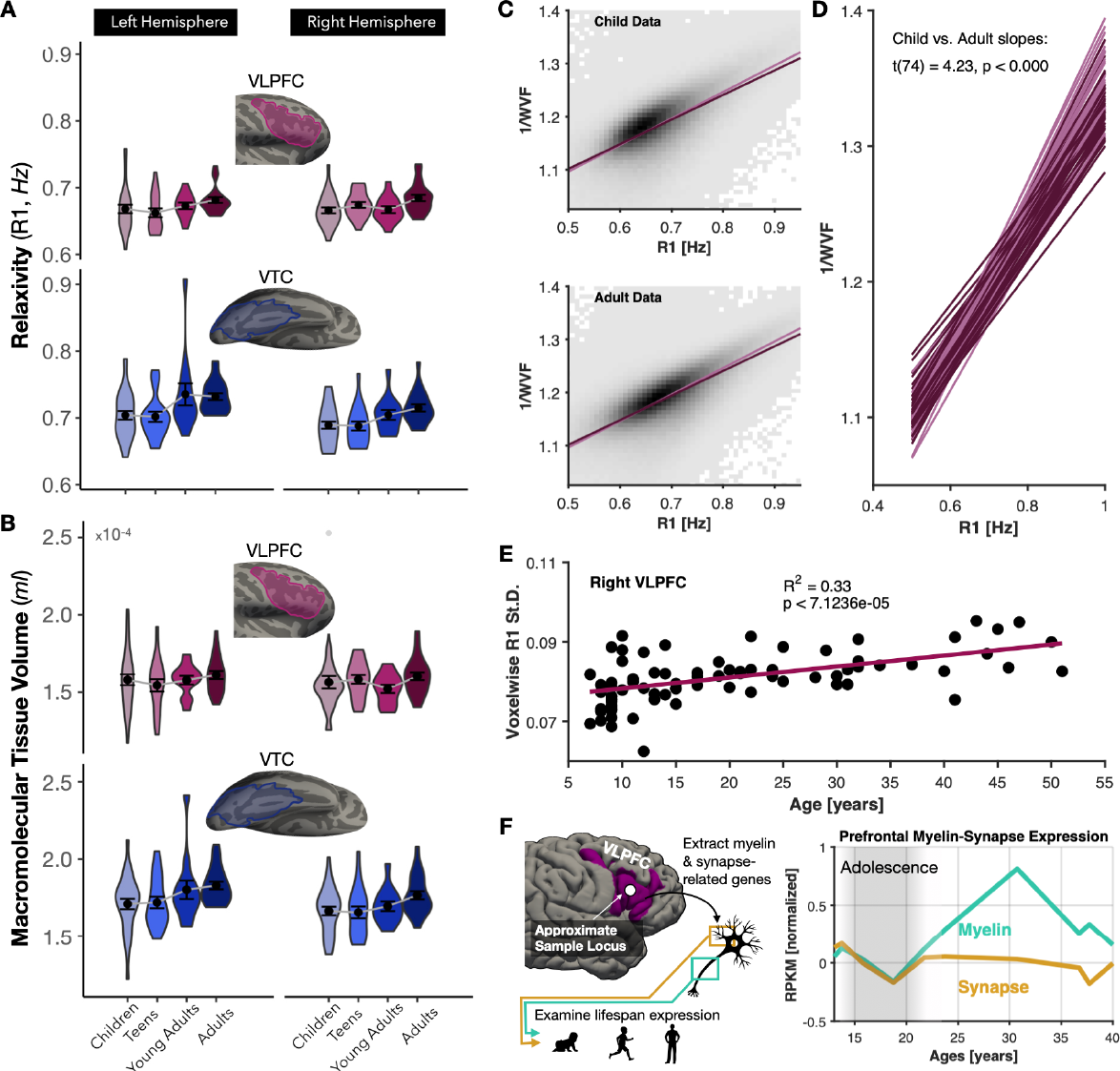
Fine-scale tissue development unfolds uniquely in prefrontal compared to temporal cortex. **(A)** Violin plots of relaxivity (R1) values for children (ages 5 -10), teens (ages 11 - 17), young adults (ages 18 - 24, blue), and adults (ages 25 - 54) in the left hemisphere (left column) and right hemisphere (right column) of VLPFC (top) and VTC (bottom). Inset cortical surfaces show example outlines of the regions over which tissue values were averaged in each participant. **(B)** Violin plots of macromolecular tissue volumes (MTV) in the same participants. Violin plots colored by age group. **(C)** Scatterplot of each participant’s age and the standard deviation of R1 values across the VLPFC ROI. Cyan line is a linear fit, R-squared and significance of fit are displayed in inset text. **(D)** Binned scatterplots of voxelwise R1 and inverse water volume fraction, e.g. the fraction of a voxel belonging to water. Voxels are pooled across all child VLPFC ROI’s (upper scatterplot) or all adult VLPFC ROI’s (lower scatterplot). The linear fit for children (light purple) and adults (dark purple) between R1 and 1/WVF are plotted in each scatterplot for comparison. **(E)** Line plots of the linear fit relating R1 and 1/WVF in voxels of each individual participant’s VLPFC ROI. Two-samples t-test comparing the slopes between children and adult lines shown in inset text. **(F)** Gene expression data of genes associated with synaptic (gold) or myelin-related (cyan) structures, averaged over all related genes within each set. Samples were extracted from the ventrolateral prefrontal cortex (tissue location shown on inset brain, left) and samples were included from donors of age 11 to 40 years old. Expression data reported in reads per kilobase of exon per million (RPKM). Data are normalized to the mean expression level during adolescence (gray shaded region, 11<age<21). Pre-normalized RPKM values for myelin-associated genes range between 9.5 and 10 RPKM, and for synapse-related genes they were between 8.2 and 8.6 RPKM.

To get a better sense of how the physicochemical environment of prefrontal cortex is changing during this period, we can further analyze qMRI metrics. There is typically a linear relationship between R1 and MTV whereby increasing MTV yields increasing R1. Deviations or changes in this linear relationship suggest the presence of hydrophobic materials that nonlinearly alter proton relaxation. That is, voxels with identical MTV values can have distinct R1 values if the physicochemical composition of the tissue is not identical. Comparing the relationship between MTV and R1 in voxels pooled from child versus adult VLPFC allows us to ask how development has sculpted the the prefrontal tissue environment. While both children and adults at a qualitative level show the expected relationship between R1 and tissue volume (plotted as inverse water volume fraction[56]), the childhood relationship appears steeper and more constrained along each axis (**Fig 4C**). When fitting this linear relationship per subject and comparing fits, we find that children indeed show significantly higher slopes (**Fig 4D**). The result of this development seems to lead to greater variegation of tissue content across cortex. We can thus ask if changing functional representations in the right hemisphere likewise change the topology of fine-scale tissue structures. Quantifying the inter-voxel standard deviation across VLPFC, we find a steady, linear change of R1 standard deviation with development, with mature cortex demonstrating more heterogeneity in its R1 landscape compared to children (**Fig 4E**). Overall, the developmental restructuring of fine-scale tissue properties in right prefrontal cortex leads to a physicochemical environment in adulthood that is distinct from childhood. When comparing two voxels, a given difference in tissue volume in adults yields a larger-than-expect difference in R1, suggesting that the adulthood cortical tissue environment is relatively biased towards tissue structures like myelin.

This suggests that the adolescent pruning of tissue we observed may have disproportionately excised non-myelinated tissue structures. We can verify this qMRI observation by examining gene expression in the same regions of human cortex across development. While RNA sequencing is not perfectly correlated to actual protein/tissue content[57] —the latter of which is measured by qMRI—we can ask if gene expression for myelin-associated genes remains relatively high following adolescence compared to non-myelinated structures such as synapses (Methods). Extracting samples from ventrolateral prefrontal cortex from a developmental gene expression dataset, we identify genes associated with both myelin structures (such as myelin basic protein) and synapses[58] and examine how their expression content varies from adolescence into adulthood (**Fig 4F**). Firstly, the gene sets associated with both myelin and synaptic structures qualitatively show a decline during the adolescent phase in ventrolateral prefrontal cortex. Following this period, we find that expression levels of genes associated with myelin rise again into adulthood while those genes associated with synaptic structures remain at their post-adolescent pruning levels. While expression of both gene sets remains well above zero, their levels relative to adolescence is consistent with qMRI metrics which suggest that development altered the physicochemical environment such that in adulthood myelin is relatively more abundant compared to non-myelinated structures like synapses or dendrites.

## Discussion

The idea that boundaries of functional representations are predictable from cortical folding was largely thought to be true only in primary sensory representations, where the boundaries of topographic projections from the thalamus conform to certain folds, such as the calcarine sulcus in vision[59,60] or Heschl’s gyrus in audition[61,62]. Recent work has overturned this notion, demonstrating that higher-level representations in occipitotemporal cortex show coupling with cortical structure[10,18-20,22,26,27,29]. Here, we extend the phenomenon of structural-functional coupling into even higher-level representations of VLPFC. In the right hemisphere, we find that face-selective responses emerge during development in the IFS between the junction with the precentral sulcus (PCS) and the first pleat of the IFS, with a second potential patch of selectivity between the second and third IFS pleats (**Figs 2-3**). The first of these face-selective regions is consistent with previous observations of face-selectivity in PFC[17,30]. Here we specify that the locus of this activity between two sulcal landmarks of the PCS and the first IFS pleat and demonstrate that this face-selective region is largely not observable until adulthood, despite clear face-selectivity posteriorly in the fusiform gyrus in the same children. We also find evidence in children and adults for an additional face-selective region in the inferior precentral sulcus just superior to the IFJ (**Fig 3**). In adults, we observe an additional region near the FEFs at the intersection of the PCS and the superior frontal sulcus (**Figs 2-3**). This constellation of face-selective regions extending from the FEF, down the PCS, and into the IFS is mirrored in the left hemisphere where a similar string of word-selective peaks can be observed.

First described nearly 200 years ago by Louis Pierre Gratiolet within non-human primates, the pleating of the cortical sheet within a larger sulcus which produces interlocking ridges and valleys was given the name *plis de passage*[46]. This was later translated to annectant gyri given that these protrusions give the illusion of connecting either side of a sulcus. While work has been done to show that many larger sulci exhibit this pleating phenomenon[47,63], whether these pleats are consistent in location and number across individuals is less clear. Given their potentially formative role in the generation of sulci during gestation[24], it is likely these folding features are under strong genetic control and might be predictors of functional boundaries. Furthermore, it is likely that this triplet pleating pattern within the IFS is human-specific[35], with findings showing this region of prefrontal cortex shows evolutionary expansion and folding patterns distinct even from chimpanzees[64]. It is thus likely these pleats are not shared with other mammalian species and represent human-unique neuroanatomy. Our structural and functional data of the IFS align with the framework that phylogenetically novel cortical regions of the human brain show the greatest ontogenetic change during childhood[65]. We find that cortical pleating within the IFS may be a developmental process that extends well beyond gestation, given that anterior portions of the right IFS may still be undergoing subtle changes in folding through childhood (**Fig 1E**), alongside its unique fine-scale tissue changes (**Fig 4**).

In VTC, functional development is primarily hallmarked by largely adult-like patterns at the age of five years old which sharpen into adulthood[2,3]. Whether or not this pattern of childhood development is a universal principle of other visually-responsive cortical regions was unknown. The developmental dynamics of VLPFC observed here, with combinations of both reductions and enhancements of responses to visual categories leading to a fundamental restructuring of functional topography, is quite distinct from the developmental scaling of childhood patterns observed in ventral temporal cortex. One possible interpretation of this effect is that VLPFC shows a uniquely protracted developmental period in which functional restructuring occurs in adolescence, while in occipitotemporal cortex such restructuring may have occurred during very early childhood. There are several previous findings in visual cortex to support this hypothesis. In VTC, developmental timepoints earlier than those studied here show pruning of representations for objects, and this pruning may extend even longer for stimuli like limbs[12,14,66]. Second, models of cortical tissue development postulate that early developmental periods display elevated synapse density followed by synaptic pruning[38,67], and in higher-level regions of the temporal or frontal lobes synapse pruning is greatly protracted and accompanied by synaptogenesis as well[39]. The qMRI data in right VLPFC with its non-linear trajectory hallmarked by post-adolescent tissue loss is consistent with such a protracted model of tissue development. When examining the physicochemical environment, we find evidence that not all tissue structures are being pruned equally, with non-myelinated structures, like synapses or somatic processes, likely being pruned to a greater extent (**Fig 4C-E**). From developmental gene expression data sampling both the same spatial region of interest (tissue near the IFS) and timepoints of interest (late childhood to adulthood), we validate our qMRI findings, showing that gene expression for myelin-related structures elevates relative to adolescence while those supporting to synaptic structures do not (**Fig 4F**).

The use here of quantitative MRI, sensitive to fine-scale tissue properties of the brain, enabled us to detect what is, to our knowledge, the first demonstration that pruning of cortical tissue microstructures can be measured in the living brain through MRI. While previous and seminal work showing reduction in gray matter volume into adulthood[68] was thought to result from pruning, newer evidence suggests this phenomenon results from myelination (e.g., tissue growth) blanching deep gray matter voxels[69], making the cortical ribbon in neuroimages appear thinner. Overall, the delayed developmental pattern in prefrontal compared to temporal cortex is consistent with recent findings showing there is extensive postnatal migration of young neurons from the subventricular zone into the frontal lobe up to a year after birth[70]. Future research should further explore the hemispheric asymmetry observed here, testing if the left hemisphere demonstrates similar functional restructuring at an earlier point in childhood compared to the right. Indeed, small but non-significant dips in left-hemisphere tissue volume (**Fig 4**) can be observed from late childhood to early teenage years in the current dataset. Auditory experiments in preterm infants (28-32 gestational weeks) found left hemisphere biases in auditory discrimination, suggesting early and strong genetic influence on left hemispheric development preceding the right[71].

Previous work has shown that PFC more consistently represents the task being performed compared to visual stimulus categories themselves[72]. That is not to say that clustered functional selectivity for visual categories like faces and words[53,73,74] does not exist, only that executive functions of VLPFC like attention or working memory impact functional topology to a larger degree than visual categories. With that said, to what extent do these developmental patterns reflect changes in bottom-up responses to visual categories, compared to top-down changes in how a task like object-based attention is represented in VLPFC? The task being performed here involved monitoring stimuli for the appearance of an oddball (a stimulus frame without an object present). Given that participants were asked to attend to the objects being presented, it is likely that the present results more closely capture changes in bottom-up representations or object-based attention. In either case, the present results support the notion that category representations in the context of oddball detection show changes in their topography from childhood to adulthood. Some of these changes (faces migrating to the right hemisphere, words to the left) are not only consistent with previous theories of lateralized visual processing[40-44,75], but extend them into the prefrontal lobe showing that it is indeed a larger phenomenon. In fact, the present results demonstrating that the relatively delayed emergence of face representations in right PFC compared to the more adult-like organization of word responses in the left suggest that for the frontal lobe, the emergence of word representations might lead that of faces. It must be stated that the exact pattern of development observed here may have appeared different if a different task had been employed, with previous work showing that the spatial extent of frontal cortex recruited during working memory shows developmental growth[76,77], with some regions such as the caudate nucleus showing decreased working memory-related activity with development[78]. Future work disentangling the development of bottom-up stimulus representations from top-down task representations will help clarify how a multimodal brain region like PFC can show multiplexed forms of functional development.

## Methods

### Participants

Functional dataset: The present analysis leveraged previously collected functional and quantitative MRI (qMRI) data[36] including 25 children ages 5 to 11 years old (M = 8.92 ± 2.18, 14 females) and 20 adults ages 22 to 26 years old (M = 23.55 ± 1.23, 15 females). All participants had normal or corrected-to-normal vision. Participants, or their parents, gave written informed consent, children assented to research, and all procedures were approved by the Stanford Internal Review Board on Human Subjects Research.

Tissue dataset: Additional analysis of cortical tissue was conducted in an independent set of participants from a large quantitative MRI dataset discussed in Yeatman et al [55]. These participants included individuals from 5 to 82 years of age (N = 109). For the purposes of our study, older adults ages 55 to 82 (N = 25) were excluded, given that the process of aging is distinct from development. Of the 84 subjects under 55, two were excluded due to missing structural data. The final 82 subjects were separated into four groups: kids ages 5 to 10 years old (N = 30, M = 8.73 ± .87, 12 females), teens ages 11 to 17 years old (N = 18, M = 13.67 ± 2.03, 10 females), young adults ages 18 to 24 years old (N = 11, M = 21.27 ± 1.74, 7 females), and adults ages 25 to 54 years old (N = 23, M = 36.70 ± 8.03, 10 females).

### Data Acquisition

Quantitative magnetic resonance imaging (qMRI): QMRI measurements were obtained using protocols from Mezer et al. [56]. For the larger qMRI dataset presented in Figure 4, four spoiled gradient echo (spoiled-GE) images were acquired to measure T1 relaxation times with flip angles of 4, 10, 20, and 30 degrees (TR = 14ms,TE = 2.4 ms, 0.8 mm x 0.8mm x 1.0 mm). To correct for field inhomogeneities, four additional spin echo inversion recovery scans (SEIR) were performed. These scans utilized an echo planar imaging (EPI) read-out, slab inversion pulse, and spectral spatial fat suppression (TR = 3.0 s, TE = minimum full value). A 2x acceleration was applied during acquisition. Inversion times of 50, 4000, 1200, and 2400 ms were used. The SEIR scans had an in-plane resolution 2.0 mm x 2.0m and a slice thickness of 4.0 mm.Using the quantitative measurements obtained from the spoiled-GE and SEIR scans, an artificial T1-weighted anatomy optimized for tissue segmentation was generated for each participant and used to reconstruct individual cortical surfaces. A second qMRI dataset was also collected using identical parameters on a separate group of participants from the larger dataset[55]. These participants were the same children and adults that completed the functional localizer discussed below. All qMRI data was analyzes with the identical mrQ analysis pipeline: https://github.com/mezera/mrQ.

Functional MRI: Participants were scanned using a General Electric Discovery MR750 3T scanner located in the Center for Cognitive and Neurobiological Imaging (CNI) at Stanford University. A phase-array 32-channel head coil was used for the category localizer experiment. Functional data were collected with a simultaneous multislice echo planar imaging (EPI) sequence and a multiplexing factor of 3 to acquire near whole-brain volumes (48 slices, TR = 1s, TE = 30ms). Slices were aligned parallel to the parieto-occipital sulcus at a resolution of 2.4mm isotropic voxels with a T2*-sensitive gradient echo sequence.

Category localizer: Participants underwent an fMRI category localizer experiment, which identifies voxels whose neural response prefers a particular category[79]. Each participant completed 3 runs, each 5 minutes and 18 seconds long. In each run, participants were presented with stimuli from five categories, each with two subcategories (faces: child, adult; bodies: whole, limbs; places: corridors, houses; objects: cars, guitars; characters: words, numbers). Images within the same category were presented in 4 s blocks at a rate of 2 Hz and were not repeated across blocks or runs. 4 s blank trials were also presented throughout a block. During a run, each category was presented eight times in counterbalanced order, with the order differing for each run. Throughout the experiment, participants fixated on a central dot and performed an oddball detection task, pressing a button when phase-scrambled images randomly appeared.

### Data Analysis

Anatomical data analysis: The SEIR and spoiled-GE scans were processed using the mrQ software package in MATLAB to produce the T1-weighted maps. The mrQ analysis pipeline corrected for RF coil bias using SEIR-EPI scans, resulting in accurate macromolecular tissue volume (MTV) and R1 (1/T1) fits across the brain. The complete analysis pipeline and its description can be found at https://github.com/mezera/mrQ. Brain-wide maps of MTV represent the volume of non-water tissue within a given voxel (1 - water fraction), while maps of R1 reflect the average rate at which protons relax to field aligned (1 / T1-relaxation time), and is more sensitive to the composition of tissue within a voxel. Voxels within regions of interest defined along the cortical ribbon were averaged within each participant’s native brain to produce a single tissue metric (average R1 or average MTV) per ROI, per subject. Regions of interest were drawn on the cortical surface, projected across the cortical depth to fill the cortical ribbon, and then brought into the qMRI data space for quantification. The VLPFC and VTC ROI’s encompass all of the sub-regions illustrated in Figure 1F-G. To reduce inter-subject noise, participants are binned within developmental age-groups: children 5-10 years old, teens 11-17, young adults 18-24, adults 25-55. From the quantitative MRI data, a synthetic T1-weighted anatomical image was produced for each individual and used for cortical surface reconstruction and functional data visualization. Artificial T1-weighted images were automatically segmented using FreeSurfer[80] (FreeSurfer 7.0.0, http://surfer.nmr.mgh.harvard.edu) and then manually validated and corrected using ITKGray to rectify any classification errors. Cortical reconstructions were generated from these segmentations using FreeSurfer 7.0.0. For Figure 1 D-E, we quantified the distance of each sulcal pleat (the pair of an annectant gyrus and its neighboring annectant sulcus) to an anatomical reference point that could be readily identified within each individual. Within each participant, we identified the inferior frontal junction as the vertex where the center of the inferior frontal sulcus’ fundus collides with the precentral sulcus. From this vertex, we measured the cortical distance to the center of the three annectant gyri within the IFS using FreeSurfer’s *mris_pmake* function.

fMRI data analysis: Functional data were motion-corrected within and between scans using the Human Connectome Project (HCP) pipeline and FSL 6.0.4. Data were then processed and analyzed using the Freesurfer Functional Analysis Stream (FSFAST) pipeline. All data were analyzed within the individual participant native brain space. Functional data were automatically aligned to the artificial T1-weighted volume produced from qMRI without smoothing, and were restricted to the cortical ribbon. Contrast maps of faces, words, bodies, objects, and places versus all other categories were produced for each participant, and where relevant, were thresholded at t-values > 2.5 in the contrast of preferred > non-preferred stimuli. That is, sub-categories (e.g., child and adult faces) were combined within a single contrast. Functional data represented in Figures 1A-B, Fig 2, and Fig 3A-C were all extracted from each participant’s native anatomical space. T-value for a given contrast map were averaged within a given ROI, hand-drawn according to anatomy in each participant’s native space. For Figure 2, the mean response within each ROI to a given category was concatenated with the mean responses to other categories individually for each group. These concatenated vectors were then compared using a Pearson correlation whose results are reported in the corner of each panel within Figure 2. For the linear discriminant analysis in Figure 3, the mean response to each category within each ROI was entered into a linear discriminant analysis using the R software package with the goal of identifying weights for each ROI-category pair that maximally separated child and adult participants. Resulting weights represent the developmental scalar that is multiplied by the mean child response to yield the average adult value. For example, a negative LDA weight applied to an initially negative child ROI-category value results in a positive adult value, signifying increase in functional selectivity to a given category within the ROI. Maps shown in Fig 3D-E were produced by using surface-based alignment to register each category contrast map from native participant space into the shared fsaverage brain space to produce average selectivity maps for each category, in each age group.

Gene expression analysis: The data reported in Figure 4F were obtained from the Allen Human Brain Atlas, in particular the BrainSpan Atlas of the Developing Human Brain and its Developmental Transcriptome. These resources are freely available at the following hyperlink: https://www.brainspan.org/. Exon microarray and associated metadata for the RNA-seq analyses summarized to genes were acquired, which samples over 17,000 genes from gestational to 40 years of age. We included data from the ventrolateral frontal cortex (VFC) region within the dataset because this tissue location was centered on the posterior inferior frontal gyrus, which is encompassed within the VLPFC ROI used within our functional and quantitative MRI data. Therefore, there is strong spatial correspondence between the gene expression and MRI data, which affords stronger comparison of these different datatypes. To identify the genes related to synaptic structures, we acquired a synaptic gene ontology set list through the SynGo database[58] (https://syngoportal.org/) which identifies, through consensus of an international research community, those genes localized to, and involved in, synaptic structures of neural tissue. For myelin-related structures, we used genes which strongly co-localize are known to be directly involved in the structure and maintenance of human myelin sheath[81]: *PLP1, PLLP, MAG, MBP, CA2, PMP22, MAL, MBP, ERMN, OMG*. Microarray expression data acquired from the human brain atlas have been preprocessed and normalized per their published pipelines. After intersecting the full developmental transcriptome gene set with our myelin and synapse gene sets of interest, we further normalized the expression of each gene to the average expression level during adolescence (11<age in years<20) within a set, before averaging across all genes within each set. Average expression levels from ages 11 to 40 years of age are then plotted in Figure 4F.

### Definitions of regions of interest

VLPFC: Ventrolateral prefrontal cortex (VLPFC) is defined as in Levy and Wagner[82] and Bugatus et al. [72]. VLPFC is posteriorly bounded by the caudal lip of the precentral sulcus and the ascending ramus of the lateral fissure (aalf). The anterior boundary lies on the most anterior edge of the pars orbitalis. The superior boundary lies on the dorsal lip of the IFS, and the inferior boundary lies on the dorsal lip of the lateral fissure. Because the frontal eye field (FEF) is heavily implicated in eye movements and viewing visual stimuli[83], we also include the inferior portion of the superior precentral sulcus, below the superior frontal sulcus, in our region of interest. An example of the VLPFC ROI can be seen in Figure 1. To achieve finer-grained analysis of the frontal lobe, we parcellated VLPFC into 9 ROIs in every individual (Fig 1F). The IFS was divided into four ROIs along its anterior-posterior axis according to annectant gyri (e.g., cortical pleats) observed consistently across participants. Each IFS subsection included one pleat: (1) the inferior frontal junction (IFJ), which extends anteriorly from the PCS to the sulcal bed of IFS-P1, (2) the posterior IFS (pIFS), which extends from the IFJ ROI to the sulcal bed of IFS-P2, (3) the middle IFS (mIFS), which begins at the anterior border of the pIFS and ends at the sulcal bed of IFS-P3, and (4) the anterior IFS (aIFS), which starts at the mIFS and extends to the end of the IFS near the lateral frontomarginal sulcus (lfms). The inferior precentral sulcus was divided into superior and inferior components (IPCS-s, IPCS-i) separated by another annectant gyrus which extends towards the IFJ. The inferior portion of the superior precentral sulcus and the surrounding gyrus were included and referred to here as the FEF ROI. We also included the op, tr, and or – which subdivide the inferior frontal gyrus – as three separate ROIs[52,84].

VTC: To define VTC, we combine cytoarchitectonic regions FG1, FG2, FG3, FG4, hOc3v, hOc4v, hO3cvla, and hO4cvlp (Fig 1G). These cytoarchitectonic regions were defined original postmortem brains, which were then resampled to cortical surface reconstructions using FreeSurfer[54]. Anatomically, we follow the boundaries proposed in Grill-Spector and Weiner 2014[53] which includes a lateral boundary at the occipitotemporal sulcus, a medial boundary at the parahippocampal gyrus, and an anterior boundary at the tip of the mid-fusiform sulcus. However, we extend VTC beyond the posterior transverse collateral sulcus, which is the posterior anatomical boundary, so that it includes cytoarchitectonic regions hOc3v, hOc4v, hO3cvla, and hO4cvlp. In our subsequent finer-grained analysis comparing VTC and VLPFC, this allows for comparison using a similar number of smaller ROIs within each lobe. Furthermore, cytoarchitectonic regions largely follow cortical folding patterns (Fig 1F), making them further comparable to the ROI’s drawn within VLPFC which were drawn relative to cortical folding patterns.

## Acknowledgements

This work was supported by startup research funds provided to JG from Princeton University. Authors JY, ZY, RM, and JG contributed to data analysis, JY and JG contributed to experimental design, and JY and JG contributed to manuscript preparation.

